# Thermoplasmonic nano-rupture of cells reveals Annexin V function in plasma membrane repair

**DOI:** 10.1101/2021.03.26.436963

**Authors:** Guillermo S. Moreno-Pescador, Dunya S. Aswad, Christoffer D. Florentsen, Azra Bahadori, Mohammad R. Arastoo, Helena Maria D. Danielsen, Anne Sofie B. Heitmann, Theresa L. Boye, Jesper Nylandsted, Lene B. Oddershede, Poul Martin Bendix

## Abstract

Maintaining the integrity of the cell plasma membrane (PM) is critical for the survival of cells. While an efficient PM repair machinery can aid survival of healthy cells by preventing influx of extracellular calcium, it can also constitute an obstacle in drug delivery and photothermal therapy. We show how nanoscopic holes can be applied to the cell surface thus allowing identification of molecular components with a pivotal role in PM repair. Cells are punctured by locally heating gold nanostructures at the cell surface which causes nano-ruptures in cellular PMs. Recruitment of annexin V near the hole is found to locally reshape the ruptured plasma membrane. Experiments using model membranes, containing recombinant annexin V, provide further biophysical insight into the ability of annexin V to reshape edges surrounding a membrane hole. The thermoplasmonic method provides a general strategy to monitor the response to nanoscopic injuries to the cell surface.

## Introduction

The eukaryotic membrane repair system involves cytoskeletal reorganization, membrane fusion events and membrane replacement.^1^ Several members of the annexin (ANXA) protein family are Ca^2+^-sensitive phospholipid binding proteins that regulate endomembrane processes including vesicle fusion, segregation and repair. ^1,2^ For example, metastatic breast cancer cells require enhanced PM repair to cope with increased frequency of injury. ^3^ To cope with such surface stresses cells respond rapidly with recruitment of several types of proteins. Components of this molecular repair kit involves several members of the annexin protein family. The ANXA2/S100A11 complex has been shown to assists in resealing the PM by enabling polymerization of cortical F-actin and excision of damaged membrane. ^3^ Moreover, ANXA7 seems to be a critical player enabling assembly of the endosomal sorting complex required for transport (ESCRT) III.^4^ Recent results also show that ANXA4 and ANXA6 act in the early phase of PM repair in cervical and breast cancer cells where they modulate wound edges by inducing curvature and constriction force, respectively. ^2^

Annexin proteins exist predominantly in the cytosol of the cell and have been found to localize to sites of membrane repair activated by influx of calcium ions during e.g. a membrane rupture.^5^ Upon binding of free Ca^2+^, annexin undergoes conformational changes, allowing it to bind to the negatively charged phospholipids abundant in the inner leaflet of the PM.^6,7^ The full scheme of the repair processes and whether they are conserved across different cellular tissues are still open questions. The specific role of annexins in membrane repair is not well understood which is mainly due to a focus on recruitment of proteins rather than the biophysical function which the proteins have on the ruptured membrane. Nevertheless, there are several proposed models of how annexins carry out cell membrane repair together with other proteins.^1^ Recently, it was shown that several annexins have the capacity to both sense and induce curvature in model membranes^2,8,9^ and also to form protein scaffolds on membranes.^8,10^ However, the implications from these biophysical effects on repair in living cells remains elusive in part due to the difficulty in performing and imaging nanoscopic ruptures. Current approaches for inducing leakage of the PM include mechanical shaking using large beads, ^11^ pulsed lasers^7^ or the use of bacterial toxins which insert into the PM and induce influx of Ca^2+^.^12^ Pulsed lasers offer an effective way to disrupt the membranes, but may cause large holes and also inflict damage to internal structures which are irradiated by the high-power pulses.

To elucidate the mechanism of action which annexins have in PM repair it is critical that nanoscopic holes can be applied to the PM in a controlled manner without extensive damage to the interior of the cells. We have developed a novel experimental assay based on thermoplasmonics to study the role of ANXAs in PM repair in live cells without compromising their viability and will allow tackling some of the open questions regarding ANXAs role in the complex PM repair machinery.

## Results and discussion

### Thermoplasmonic injury to the PM

Holes in living cells recruit a number of different membrane repair associated proteins to the site of injury. Using a combination of inorganic plasmonic nanoparticles and a highly focused *λ* = 1064 nm laser coupled to a confocal microscope, we examine the ANXA5 protein responses to PM injury.

By optically heating plasmonic nanoparticles near the PM (Fig. 1A) we form nanoscopic holes in the PM of living cells. Local puncture of the PM leads to influx of Ca^2+^ which mediates binding of ANXA5 to the PM. By using the calcium sensitive dye Fluo 4-AM we observe influx of Ca^2+^ upon irradiation of a AuNP (Supplementary Fig. 1). Irradiation of plasmonic nanoparticles provides an efficient way to perturb the membrane of cells locally and has been successfully demonstrated as a method for fusing cells and membranes.^13^

**Figure 1:**
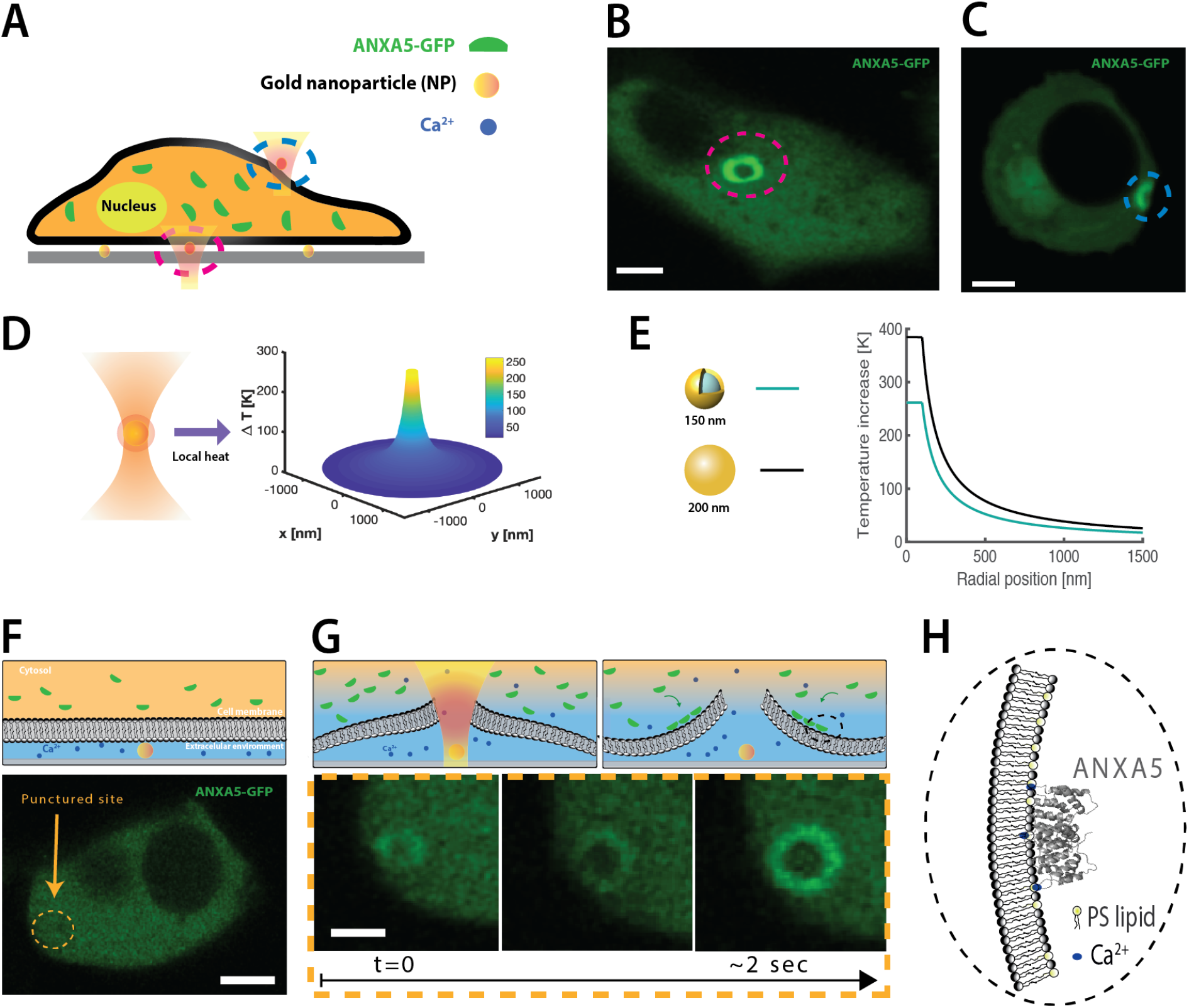
Outline of the experimental assay for applying a nanorupture to a plasma membrane. (A) Schematic illustration of a ANXA5-GFP transfected cell plated on a glass with immobilized AuNSs. (B) Fluorescence confocal image from a ANXA5-GFP transfected HEK293T cell injury from below the cell by irradiation of an immobilized AuNP. (C) Fluorescence confocal image from a ANXA5-GFP transfected HEK293T cell injury on the side of the cell using an optically trapped AuNP. (D) Heating of a AuNS with a NIR laser induces localized heating from the particle. (E) The heating profile of two different size NPs (AuNP, *d* = 200 nm and AuNS, *d* = 160 nm) used in this work at a laser intensity of *I* = 7.6 · 10^10^ W/m^2^. (F) Before the NPs are irradiated with NIR laser light, the plasma membrane is stable adhered on top of the particles. (G) After the thermoplasmonic rupture, ANXA5-GFP up-concentrates in a ring-like fashion around the wounded site (confocal image in the bottom part) in less than two seconds after irradiation. (H) On the top right side an illustration of Ca^2+^ mediated binding of ANXA5 to the PM. Crystal structure of ANXA5 from the PDB entry: 1a8a.

Visualization of the cellular response to optically triggered local PM perturbation, is carried out by parallel imaging using confocal fluorescence microscopy. To achive the best imaging of the ANXA5 response to rupture we chose to immobilize the nanoparticles on the surface underneath the cell. Near infrared (NIR) light was used to irradiate the gold nanoparticles (AuNP, *d* = 200 nm) or gold nanoshells (AuNSs, *d* = 160 nm). The AuNS are designed to be resonant with the NIR light whereas the AuNPs exhibit significant absorption in the NIR region and hence both structures facilitate high absorption and heating at the wavelength (*λ* = 1064 nm) used here (see Fig. 1D,E). The solid gold nanoparticles have been found to exhibit a higher thermal stability than the gold nanoshells, ^14,15^ but to achieve the same heating as the AuNSs their diameter has to be slightly larger.

The high temperature region surrounding irradiated NPs extends only ~ 100 nm from the nanoparticle, see Fig. 1D,E.^16^ This makes it possible to apply extremely localized ruptures to the PM which are localized in all three dimensions. This is unlike the effect from pulsed lasers that also ablate internal cell material along the path of the laser light. This allows us to test the cellular response to PM ruptures of nanoscopic dimensions in both cells and model membranes. Nanoscopic holes may be frequent during aggressive cellular migration and invasion^1,17^ and therefore investigation of protein recruitment in response to nanoscopic ruptures are relevant for understanding how living cells cope with ruptures.

The assay described in Fig. 1A was used to investigate the cellular rescue response to plasmonic injuries underneath human embryonic kidney 293T (HEK293T) cells on a glass surface. By immobilizing nanoparticles on the surface of the glass and seeding the cells on top, a two dimensional visualisation of the injury was obtained. The results show up-concentration of ANXA5 in a ring-like fashion around the wound site in HEK293T cells (see Figs. 1B,C,F and G).

To minimize the thermal effect on the cell we only briefly irradiated the NP for ~ 1 s which minimizes damages to proteins which typically need more time to unfold. The ring subsequently formed during a few seconds as shown in Figs. 1F and G.

These data are in good agreement with previously reported data of how a similar protein, ANXA4, responds to PM injuries applied using a pulsed laser source in Ref.^2^ ANXA4 was recruited to the site of injury and the curvature induction effect of ANXA4 was expected to curve the membrane away from the site of injury. The shape and structure of ANXA5 (Fig. 1H) resembles ANXA4 and we anticipate that the internal binding of ANXA5 around the holes in our experiments would generate some curvature in the membrane as schematically indicated in Figs. 1G,H.

Calculation of the ANXA ring sizes was based on the full-width-half-maximum of the intensity line profile as depicted in Fig. 2A. The resulting radii for a collection of more than 130 cells damaged via plasmonic heating of AuNPs and AuNSs are shown in Fig. 2B. Following irradiation, all injuries were observed to recruit ANXA5 to the wound site within seconds (see Figs. 1F and G), visible as a ring surrounding the wound. The ANXA5 ring size varied from hundreds of nanometers to several microns, suggesting that the PM repair machinery might be more efficient for some cells. Whether this is due to the cell cycle stage, the hole size in general or the amount of transfected ANXA5, is something that should be addressed in future experiments.

**Figure 2:**
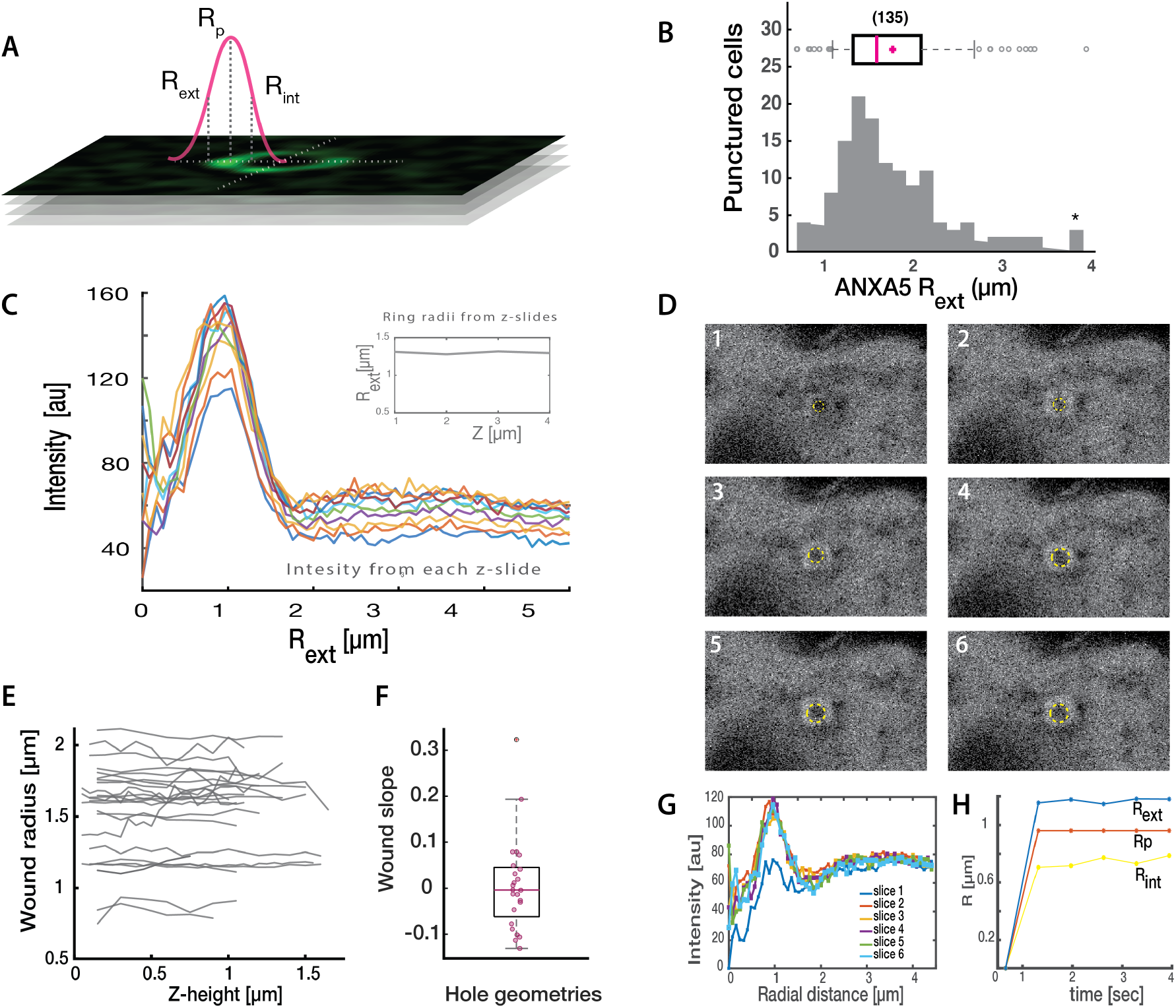
Quantification of the ANXA5-GFP ring sizes surrounding the wounded sites. (A) Schematic of how the analysis is carried out with regard to radius determination. (B) Histogram of the ANXA5 ring radii measured for all experiments. A boxplot from the histogram has been placed on to the upper part of the graph. (C) Fluorescence intensity line profiles across the ANXA5-GFP rings for each slice on of a z-stack. The inset shows the variation on the radius size along each slice of the z-stack. (D) Hole evolution over time. The time dependent hole evolution shows how ANXA5-GFP is upconcentrated immediately after injury. The wound size is stable following ANXA5-GFP upconcentration. (E) Wound radii determined by the analysis method presented in (A) as a function of wound depth. (F) Slope of ring radius versus height extracted from the analysis data in (E). (G) Fluorescence intensity line profiles across the ANXA5-GFP rings for each time frame (0,66 seconds of time interval per frame). (H) Time evolution of the ANXA5-GFP ring radii from (D) calculated from the data in (G).

Cell viability was detected by monitoring the binding of ANXA5 on internal membranes. Excessive influx of Ca^2+^ leads to translocation of ANXA5 from the cytoplasm to internal membranes including the nuclear membrane and was detected in a few cases. In these few cases cell death occurred within a few minutes and such data were not used to quantify the shape of the ANXA5 ring. The vast majority of thermoplasmonically ruptured cells showed no signs of ANXA5 binding to internal membranes, distal to the site of injury, even after 2h of irradiation which is a sign of successful membrane repair.

For several injuries, only a fraction of a ring shape was observed (See Supplementary Fig. S1 panel D and G). This phenomenon might be because the membrane disruption happens in an unfavorable way in parts of the injury, an asymmetric adhesion of the PM to the substrate or interactions with the actin cortex underlying the PM.

It has been reported that different members of the annexin family are able to roll membranes^2^ as well as sense curvature. ^8,10^ Such biophysical effects could lead to rolling of the membrane edges away from the hole and lead to a ring shaped structure if the membrane was free to roll.

To further investigate the actual three dimensional shape of the ANXA5 ring we performed confocal microscopy of each injury at several positions along the axial direction. ANXA5 rings induced by the thermoplasmonic-membrane rupture showed no change of the ring-size with height, see Fig. 2C.

The intensity profiles shown in Fig. 2C were observed in several other z-stacks and the extracted radius versus height was calculated for each injury, see Fig 2E. The procedure behind the calculation of the ring sizes was the same as shown in Fig. 2A, keeping the same center coordinates for the ring in all images within the z-stack. From the analysis of the radii of each z-stack (see Fig. 2E), the slope of the resulting radius profile was calculated. A slope of zero value would indicate no change in the radius of the ANXA5-ring along the z-direction.

As shown in Fig. 2F the slopes of radius versus height was close to zero in all cases. This strongly suggests that under these conditions the ANXA5 does not significantly bend the cell surface towards the cytoplasm, but rather forms a cluster of ANXA5 surrounding the site of injury. To test for a possible presence of a membrane roll resulting from ANXA5 binding to the PM we imaged the time evolution of the ring formation. As shown in Fig. 2D,G,H the ring reaches its final radius within ~ 1 s and within the time resolution of a confocal microscope we see no gradual increase in the width of the ring which is expected to be the case if the PM was rolling away from the site of injury.

To gain further insight into the response of cells when punctured at a location which is not in contact with an adhesive substrate, we also punctured cells from the side using plasmonic nanoparticles optically trapped in 3D. Such experiments also show an upconcentration of ANXA5 around the site of injury, see Fig. 1C and Supplementary Fig. S6. Interestingly, these experiments occasionally showed a slight inward bending of the area enriched in ANXA5 as shown in Fig. 1C.

To isolate the mechanistic effect of ANXA5 on the membrane surrounding the rupture we used Giant Unilamellar Vesicles (GUVs) encapsulating recombinant annexins as a minimal model system. GUVs can be readily punctured using thermoplasmonics which has been utilized for vesicle fusion^13^ and permeability studies of model membranes. ^18^ Following the same experimental approach to generate punctures we encapsulated recombinant ANXA5-GFP or the cognate protein ANXA4-GFP inside GUVs by using the polyvinyl alcohol (PVA) gel hydration method (see Supplementary Materials). Briefly, a PVA gel coated glass slide is covered with the desired lipid mixture and incubated with a protein-containing growing buffer for GUV formation followed by isolation of the GUVs in an observation buffer.

The presence of millimolar concentrations of Ca^2+^ ions in the observation buffer facilitates the binding of ANXA5 to the internal membrane due to residual leakage of ions across the membrane (See Figure 3). A total number of N=11 GUVs were punctured and two very distinct effects were observed repeatedly upon membrane rupture: i) more severe ruptures using *P* ≈ 120mW and applied to GUVs encapsulating ANXA4 or ANXA5, respectively, resulted in expansion of the hole and with an increasing membrane roll forming at the rim of the hole as shown in Fig. 3D. Moreover, using recombinant ANXA4 we were able to replicate ring-like upconcentration scaffolds in GUVs - see Fig. 3C. ii) Under moderate heating exposure of *P* ≈ 50mW, small pores were formed and closed after a short time and we observed invaginating buds of the membrane forming inside the vesicle (Fig. 3B). There was a remarkable up-concentration of ANXA5 in the neck region of this bud-like structure created after membrane rupture - see yellow circled area in Fig. 3B which is consistent with the curvature recruitment reported for ANXA5.^10^ The buds were not successfully released from the membrane after wound closure. Application of small injuries several times at the same location, resulted in accumulation of inward pointing buds before vesicle collapse (Fig. 3E).

**Figure 3:**
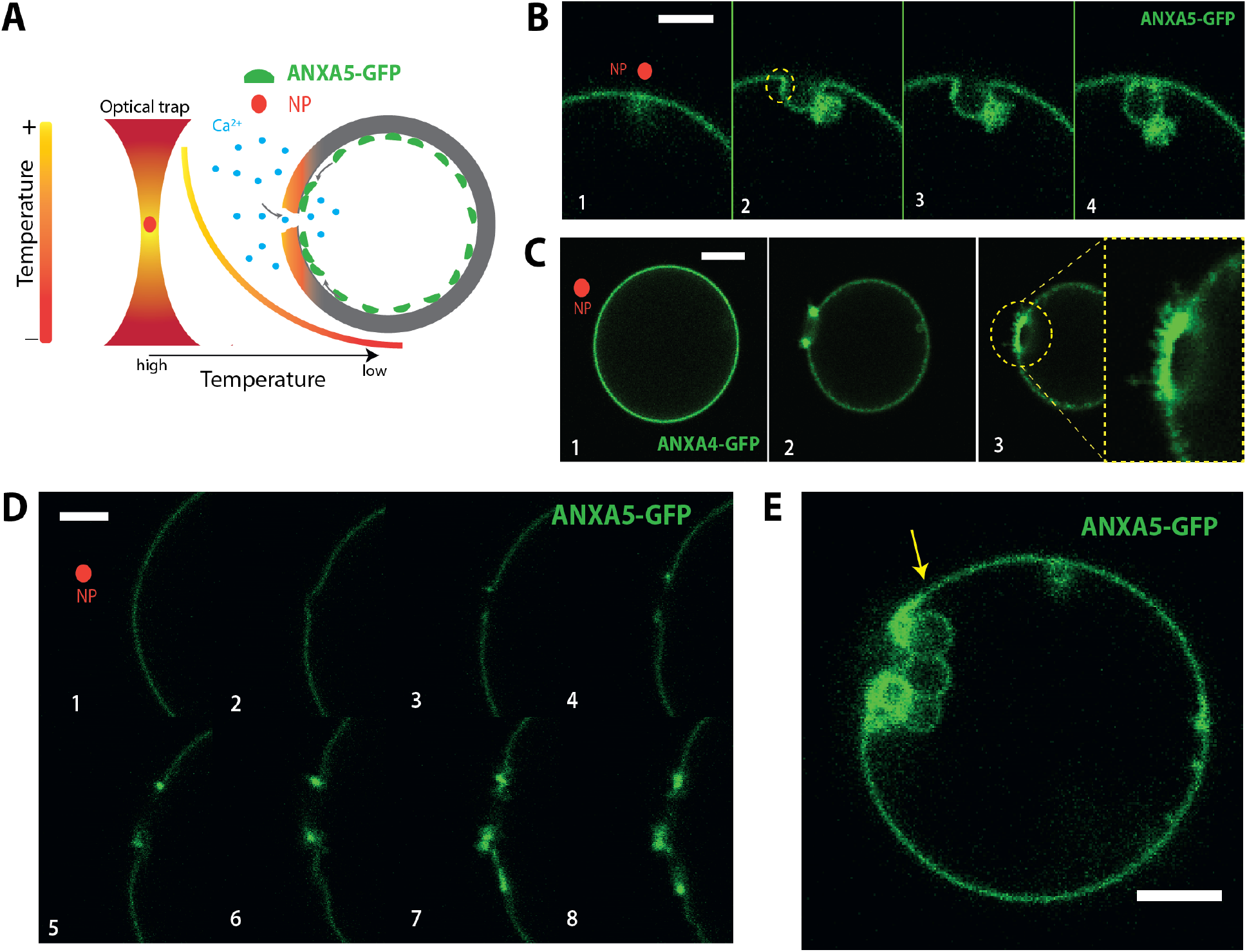
Plasmonic rupture in GUVs with encapsulated GFP-ANXA5. (A) Schematic illustration of the experiment with ANXAX5 encapsulated in a GUV. A trapped AuNS in the vicinity of the GUV will generate enough heat to provoke rupturing of the membrane when sufficiently close. (B) Montage with the time evolution of a hole created in GUV with ANXA5 bound to the membrane. Scale bar is 2*μm*. (C) Montage with the time evolution of a GUV rupture with the cognate ANXA4 protein bound to the membrane. Scale bar is 10 *μm*. (D) Montage with the time evolution of a GUV rupture with ANXA5 bound to the membrane. Scale bar is 5*μm*. (E) GUV after repeated thermoplasmonic puncturing. Scale bar is 10 *μm*.

These experiments embody very well the different effects that ANXAs have on pure membranes where no interactions with the cytoskeleton are present. The apparent rolling of membranes away from the site of injury shown in Fig. 3C,D and also for the membrane signal in Supplementary Fig. S5, is a signature of the curvature induction by ANXA4 and A5 reported in Refs.^2,8^ Additionally, accumulation of ANXA5 was observed at the neck regions connecting the spherical bud to the GUV membrane which is a clear signature of membrane curvature sensing by ANXA5 (yellow circle in Fig. 3B). The mechanism behind the budding events is unknown, but could result from ANXA5 induced inward bending of the membrane surrounding the hole into a funnel shape, followed by resealing of the funnel and subsequent formation of the spherical bud. Repeated injuries resulted in a several buds which were not released into the GUV lumen, but rather clustered near the site of injury Fig. 3E.

Several annexins have been shown to aggregate membrane vesicles and hence they have the ability to bridge or cluster apposing membranes as shown in Fig. 3E. ANXA5 isolated from chicken^19 20^ and human ANXA2^21^ have been reported to mediate contact between apposing membranes through interactions between the N-terminals in the proteins. We have also observed the same interaction in human ANXA5. When recombinant ANXA5 was added in a 2mM Ca^2+^ buffer it would bind to the GUV’s membrane and in time aggregate the vesicles, see SI Fig. S3. Such interactions could well play a role in both the vesicle clustering and rolling behavior observed in Fig. 3.

Our experiments *ex-vivo* and in model membranes show that ANXA5 mainly cluster around the site of injury in a ring like form. It is difficult from the current data to conclude whether the extensive rolling observed in model membranes (Fig. 3 and in Ref. ^2^) also takes place in cells. Cells do indeed have an actin cortex underlying the PM which could inhibit the efficient rolling observed in model membranes exposed to ANXA5. Instead, we propose a more complex model of how ANXA5 could stabilize the PM around the wound area based on the results presented here - See Fig. 4B. Initial rolling will be counteracted by the presence of actin network, but the PM could be allowed to fold back onto itself by two ANXA5 proteins interacting via their N-teminals as shown in Supplementary Fig. S3. Morever, extensive binding of ANXA5 to the PM surounding the hole is known to result in a connected protein scaffold^7^ and could counteract expansion of the hole. This would allow the cell to repair the hole through other mechanisms like clustering and fusion of internal vesicles to the ruptured membrane. Such a mechanism is in good agreement with the heterogeneity of the ANXA5-ring we found in our experiments.

**Figure 4:**
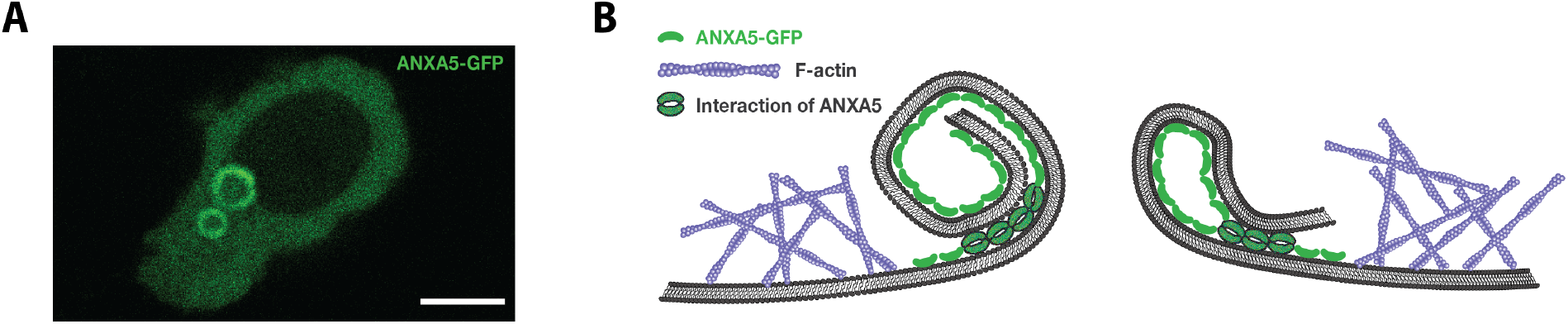
Proposed model for the structure of ANXA5 at a wounded site. (A) Confocal fluorescence images from thermoplasmonic experiment in ANXA5 transfected cells with two puncture sites. Scale bar is 10 *μm*. (B) Binding of ANXA5 to free edges induces local bending and interaction of ANXA5 molecules with the cytoskeleton arrests the membrane rolling deformation and aids to stabilize the would.

## Conclusion

Cells show rapid recruitment of ANXA5 in response to membrane ruptures induced by thermoplasmonics. Upon irradiation of plasmonic gold nanostructures located on the plasma membrane, Ca^2+^ leaks into the cytoplasm and triggers ANXA5 binding to the internal PM. ANXA5 is found to form long lasting ring-like structures around the site of injury within few seconds following rupture of the PM. The fluorescent signal from the ring could arise from folding of the ANXA5 bound membrane caused by the curvature generation by the protein. The use of plasmonic puncturing of the cell surface combined with confocal imaging offers an excellent platform for identifying molecular components involved in surface repair. Considering the efficient uptake of nanoparticles in cells we envision that the precision provided by this approach will allow us to resolve the mechanisms employed by cells to heal internal damage in nuclear or endosomal membranes.

## Experimental

### Cell culture

HEK293T cells (ATCC) were cultured in Dulbecco’s modified Eagle’s medium supplemented with 10% fetal bovine serum (FBS) and 100 units/mL penicillin and 100 *μ*g/mL streptomycin (Fisher). The cells were maintained in a humidified incubator with 5% CO_2_ at 37°C. Expression plasmid with turbo-GFP C-terminal tag containing human ANXA5 cDNA were purchased from OriGene Technologies. HEK293T cells were transiently transfected with the indicated plasmid using Lipofectamine LTX transfection reagent (Fischer) according to the manufacturers protocol 24 h before they were used in experiments

### Reagents

Gold nanoparticle colloid suspension was created by adding 1 *μ*L AuNPs (100 nm or 200 nm) from stock solution to 100 *μ*L observation buffer and sonicating for 2-5 min before use. The AuNS are coated with Poly(ethylene glycol) (PEG) in order to prevent aggregation of the particles and stabilize them in solution. Gold nanoshells (AuNS) were also used, their core consisting of a 120±4*nm* silica sphere upon which a thin gold shell (20±7*nm*) has been grown giving the complex a total diameter of ~ 160nm (Nanocomposix, CA USA). To generate C-terminal GFP-tagged recombinant Annexin A5 (ANXA5) protein, ANXA5 cDNA was subcloned into pETM11SUMO3sfGFP.^22^ Proteins were produced using Immobilized Metal-Affinity Chromatography (IMAC) followed by fast protein liquid chromatography (FPLC).

### Surface coating with particles

The nanoparticles used for the experiments were prepared a day prior to imaging. First an aliquot of particles was sonicated for 20-30 minutes to make sure they did not aggregate. Poly-L-Lysine (PLL) (Gibco) was used to make the particles and cells stick better to the surface of the microwell (MatTek 35mm glass bottom dish). 500*μ*l PLL was added to the microwell and incubated for 10 minutes. The well was washed with distilled water (Gibco) and dried inside a flowhood. When the microwell was dry, the particles were added to the microwell with 500*μ*l of DMEM.

### Optical trapping and imaging

Confocal imaging was performed on a Leica SP5 confocal microscope into which an optical trap, based on a 1064 nm laser (Spectra Physics J201-BL-106C), was implemented. ^23^ Optical trapping was done at the focal plane of the microscope and by using a Leica PL APO, NA=1.2, 63X water immersion objective to tightly focus the laser light. The optical trap was stationary, but the trap could be moved relative to the GUVs by translating the sample which was mounted on a piezoelectric stage (PI 731.20, Physik Instrumente, Germany) allowing lateral movements with nanometer precision. A glass bottom open chamber or microwell, containing either the GUVs or HEK293T cells, molecular fluorescence probes and gold nanoparticles, was mounted on the microscope and kept at room temperature during the experiment. The 488 nm argon laser line was used to excite the GFP fluorophores. To detect the scattered and reflected light from the AuNPs, a 613 nm or 476 nm laser line was used in reflection mode.

### Plasmonic heating

The temperature profile around an irradiated NP can be found by solving the heat transfer equation^24^ where the local heat intensity comes from light dissipation inside the irradiated NP,^25^ the temperature increase Δ*T* (*r*) can be written as a function of distance to the source (*r*) as:^26^

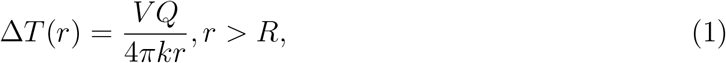

where *R* is the radius of the spherical NP, *k* is the thermal conductivity of water (0.58 W/mK in water at room temperature), *V* is the volume of the particle and *Q* is the local heat dissipation.^27^ From Ref.,^28^ equation (1) can be re-written to relate the increase of temperature to the cross section absorption *C_abs_* as

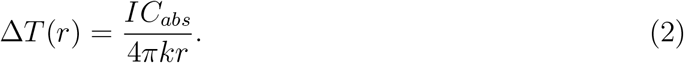

Δ*T* (*r*) is the temperature increase relative to ambient temperature and *κ* is the thermal conductivity. *Q* = *IC_abs_* is a measure of the generated heat (the amount of heat produced per unit time and volume inside the NP), largely contributed by Joule heating inside the NP. *I* is the intensity of the laser irradiation at the sample plane. The temperature profile around an irradiated NP is governed by eq. 1, which reaches a steady state within tens of nanoseconds, satisfying Laplace’s equation. The solution for the temperature gradient will be a simple function of distance, *r*, to the surface of the NP with radius of *R*.

Through Mie based calculations, ^29^ we can calculate the absorption cross section, *C_abs_*, from which the temperature profiles of irradiated strongly absorbing nanoparticles^30^ can be obtained. Fig. 1B shows equation 2 plotted as a function of distance to the particle center.

### Vesicle swelling

Polyvinyl alcohol gel (PVA) (5%, 50 mM sucrose, 25 mM NaCl/Tris) was heated at 60°C for 20 min. Glass slides was first cleaned in ethanol and dried with ultra spray, followed by heating in an air Plasma Cleaner model Harrick Plasma cleaner PDC-002. 90 *μ*L of warm PVA gel was applied on the clean glass slides, after which they were heated in a 50°C oven for 50 min. Subsequently, the vesicles were grown on top of the prepared PVA glass slides. A 50 *μ*L glass Hamilton syringe was cleaned in chloroform to ensure minimum contamination. 30 *μ*L of prepared lipid mix (95% DOPC, 5% DOPS) was added on top of the PVA gel layer on the glass slides. The applied lipid mix was dried with nitrogen for 30 seconds followed by vacuum drying for 2 hours. A clean vesicle chamber was used to grow the vesicles. 350 *μ*L growing buffer (NaCl 70mM, Tris (pH 7.4) 25 mM, Sucrose 80 mM) with possible recombinant annexin was added to the chamber on top of the PVA-coated glass slide and the vesicles were left to form for ~1 hour. 340 *μ*L of vesicle and buffer solution was collected from the chamber and deposited in an Eppendorf tube. 300 *μ*L observation buffer (NaCl 70 mM, Tris (pH 7.4) 50 mM, Glucose 55 mM) were added for the vesicles to sink to the bottom.

### Protein encapsulation

Annexin A5 (900nM) was added to the prepared cover glass. Followed by an addition of the prepared growing buffer at room temperature (total volume of 300 L) was then set to form encapsulated annexin A5 giant unilamellar vesicles for ~1 hour. The vesicle solution was extracted by pipetting directly from the cover glass into an Eppendorf tube and 1000 *μ*l observation buffer. The vesicles were set to sink for 5 minutes followed by centrifugation at 600 rcf for 10 minutes at 13°C. Subsequently the added observation buffer was carefully removed from the top and down to ensure minimal loss of GUVs and to remove excess annexin A5 on the outside.

### Data analysis

All images were analyzed using Matlab (The MathWorks, Inc., Natick, Massachusetts, United States) as well as the temperatures calculated in Fig. 1D and 1E were plotted using Matlab and calculations were based on Mies equations. ^29^ Fig. 2 were calculated using an in-house Matlab workflow that is available on request. All confocal microscopy images were processed using Fiji ImageJ distribution.^31,32^ Additionaly, for supplementary figures S1 to S3, S5 and S6 FigureJ plugin^33^ was used.

## Acknowledgement

This work is financially supported by Danish Council for Independent Research, Natural Sciences (DFF-4181-00196), by a Novo Nordisk Foundation Interdisciplinary Synergy Program 2018 (NNF18OC0034936), by the Scientific Committee Danish Cancer Society (R90-A5847-14-S2), the Lundbeck Foundation (R218-2016-534) and a Lundbeck Foundation Center of Excellence (Biomembranes in Nanomedicine).

