## Supplementary Information for "Thermoplasmonic nano-rupture of cells reveals Annexin V function in plasma membrane repair"

### Supporting Information Available

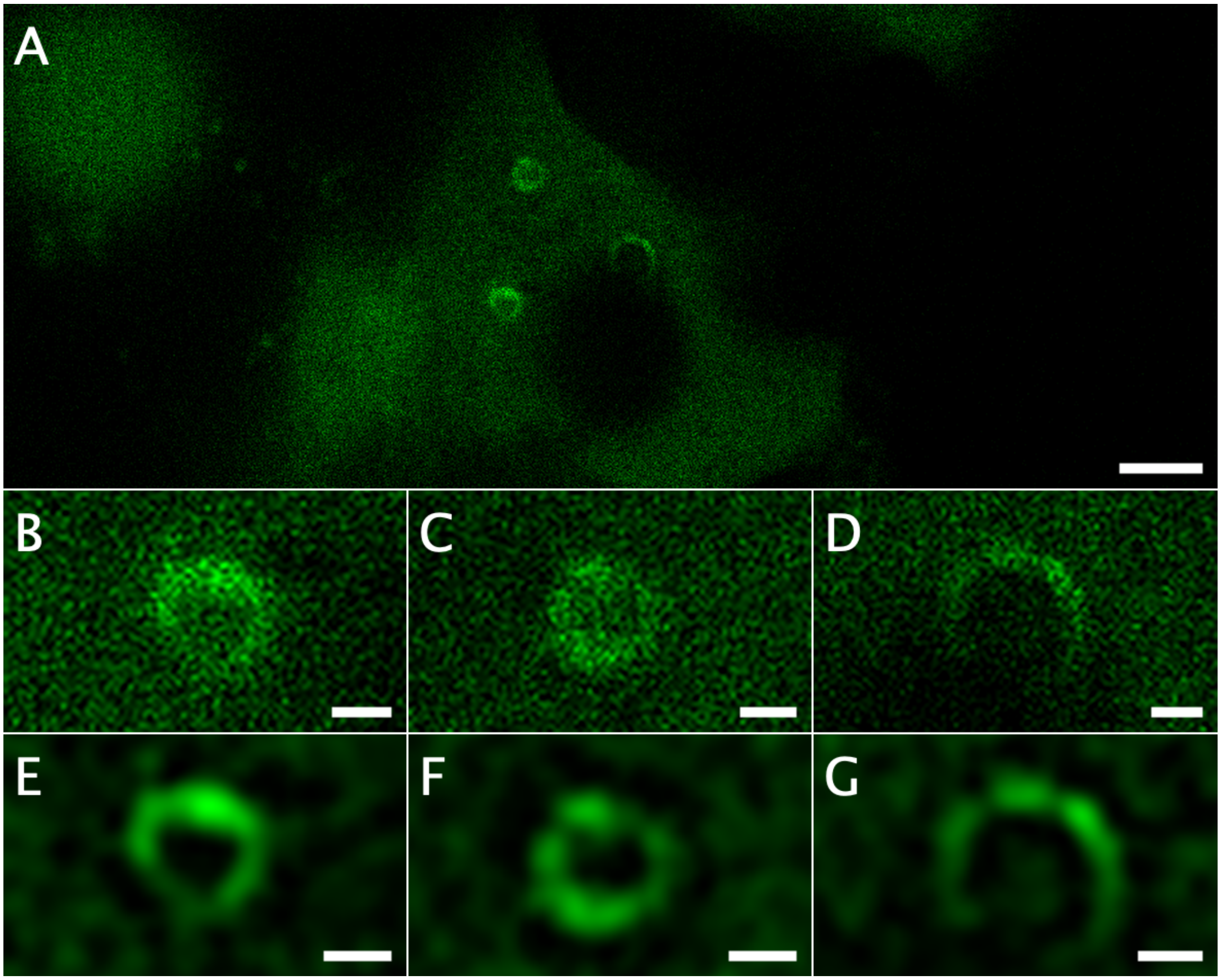


**Fig. S1.** Membrane puncturing of cell. The cell is able to survive puncturing several times by activation of the PMR machinery. A) Cell with 3 punctures. Scalebar is 5 microns B), C) and D,) show the unedited rings from A. E), F) and G) show edited images using a Difference of Gaussian (DoG) filter of the images above. All scalebars, except for A, are 2 $\mu m$.

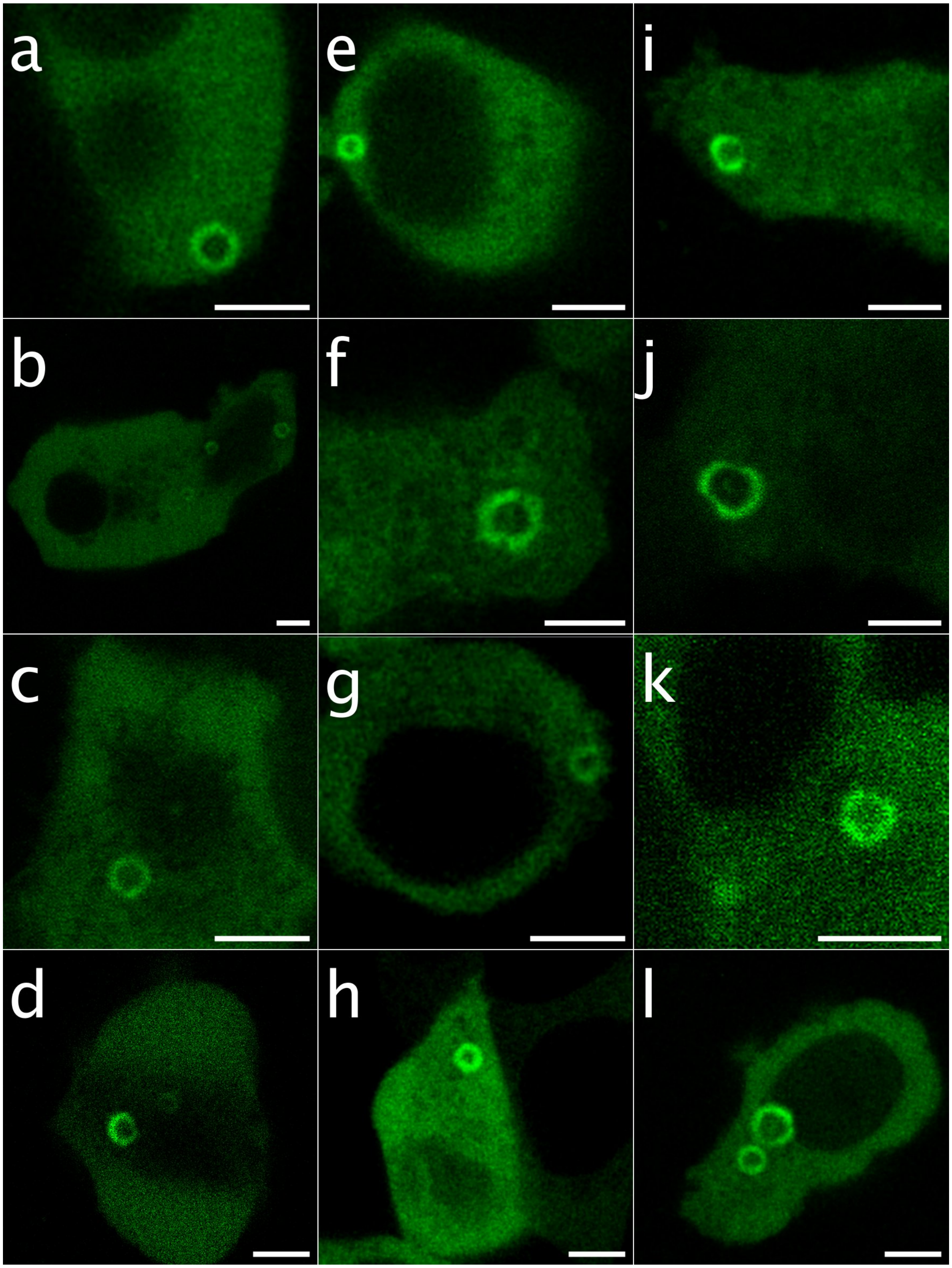


**Fig. S2.** . Example of punctured cells expressing ANXA5-gfp. Scale bars are all 5 $\mu m$.

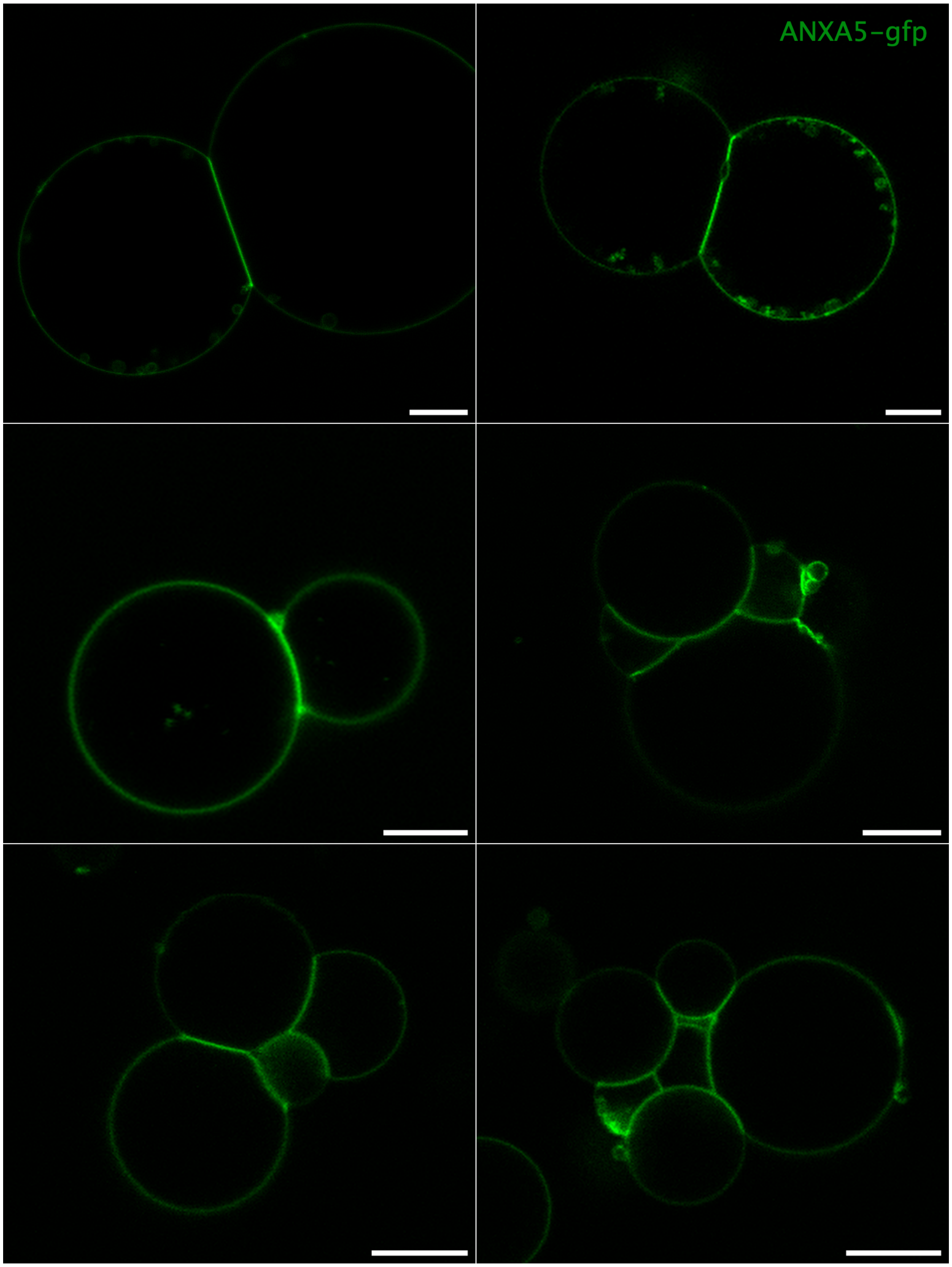


**Fig. S3.** . Interaction of C-terminal in between ANXA5.These aare examples of GUV prepared using electroformation (DOPC 90% + DOPS 10%) in a CA2+ buffer of 2mM. Purified ANXA5-gfp was added in solution causing strong binding of GUVS. Scale bars are all 10 $\mu m$.

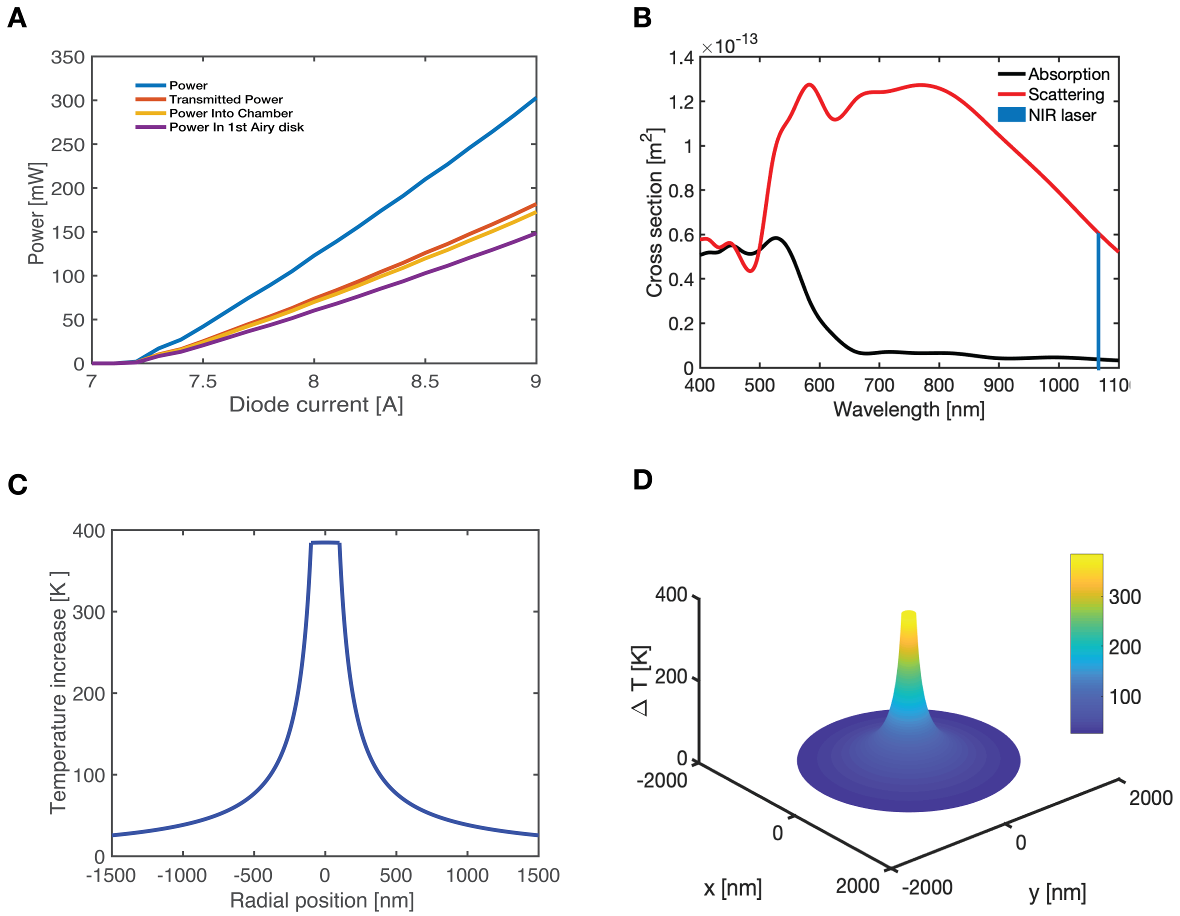


**Fig. S4.** . Gold nano-particle 200 $nm$ calculations for thermoplasmonic heating. (A) Power measurements of the experimental set up at different points of the NIR laser path. Blue, right after the fiber. Red and orange are before and after the objective respectively. Purple is the power measured in a sample. (B) Absorption and scattering cross sections calculated for a AuNP of 200 nm diameter. (C) Using a 8 A current through the laser diode can calculate using (A) and (B), the heat produced by the AuNP. (D) 3D render of the same heating profile presented in (C).

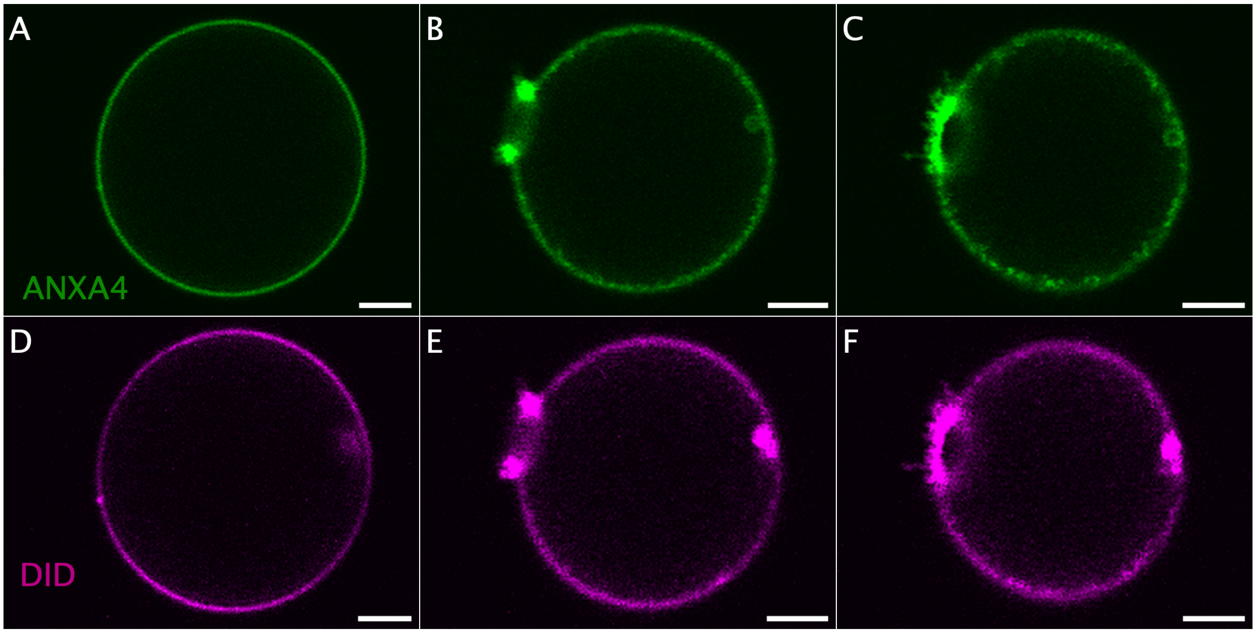


**Fig. S5.** . Example of punctured GUV with bound ANXA4-gfp and bilayer DiD labeling. Scale bars are all 5 $\mu m$.

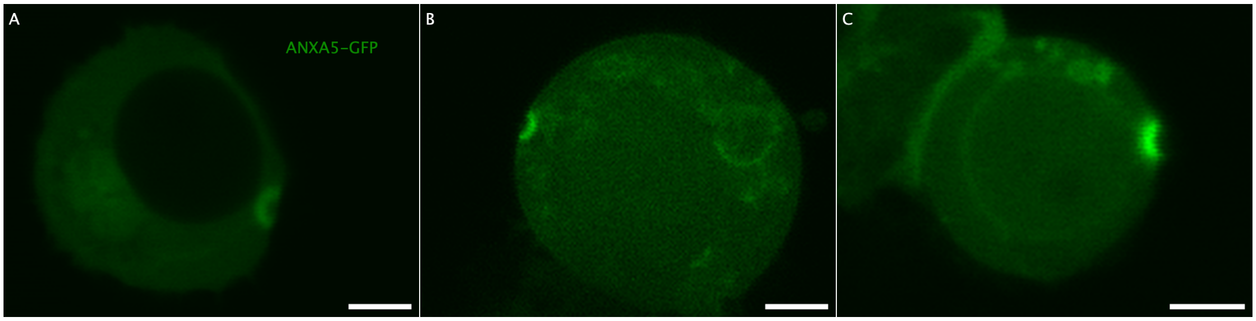


**Fig. S6.** . HEK293T cells expressing ANXA5 punctured from one side. Scale bars are all 5 $\mu m$.
